# Reproducibly oriented cell divisions pattern the first flat body structures to set up dorsoventrality and de novo meristem formation in *Marchantia polymorpha*

**DOI:** 10.1101/2024.07.08.602509

**Authors:** Eva-Sophie Wallner, Liam Dolan

## Abstract

Land plant bodies develop from stem cells located in meristems. However, we know little about how meristems initiate from non-meristematic cells. The haploid body of bryophytes develops from unicellular spores in isolation from the parental plant, which allows all stages of development to be observed. We discovered that the Marchantia spore undergoes a series of precisely oriented cell divisions to generate a flat prothallus on which a meristem later develops de novo. The young sporeling comprises an early cell mass. One cell of the early cell mass elongates and undergoes a formative division that produces the prothalloblast, which initiates prothallus formation. A symmetric division of the prothalloblast followed by two transverse divisions generate a four-celled plate that expands into a flat disc through oblique divisions in three of the four plate cell-derived quadrants. One quadrant gives rise to a flat flabellum. A notch with a meristem and apical stem cell develops at the margin of the flabellum. The transcription factor MpC3HDZ is a marker of the first flat prothallus structure and polarises to the dorsal tissues of flabella and meristems. Mp*c3hdz* mutants are defective in setting up dorsoventrality and thallus body flatness. We report how a regular set of cell divisions forms the prothallus – the first dorsoventral structure – and how cells on the margin of the prothallus develop a dorsoventralised meristem de novo.

## Introduction

Meristems are the morphogenetic centres containing stem cell niches that produce most tissues and organs of multicellular land plant bodies. In the diploid phase of vascular plant life cycles, the first meristems develop on the embryo, which in turn is derived from the zygote ensheathed in tissue ^1,2^. These meristems form the first modular axes of the metameric plant, and the initiation of new meristems throughout the life of the plant generates the adult body ^3^. Meristems also form the haploid body of the bryophyte life cycle ^4,5^. In bryophytes, meristems organise among groups of cells derived from the free-living spore cell ^6,7^. These meristems form the first modules of the plant, and like vascular plants, the initiation of new meristems generates the adult body ^7^.

On germination, the haploid spore of the liverwort *Marchantia polymorpha* forms a prothallus on which the meristem that gives rise to the entire plant body develops (Figure 1A) ^7,8^. The first step in this process is the asymmetric cell division of the spore to produce an apical germ cell and a differentiated rhizoid cell ^4,9-11^. The apical germ cell typically undergoes several rounds of divisions forming a morphologically variable cluster of cells that has been referred to as cell mass ^12^, protonema ^7^ or germ tube ^4,12^. Compared to the tube-like protonema of mosses ^13^, *Marchantia polymorpha* typically forms a tight cluster of largely spherical (isodiametric) cells when grown in standard white light conditions (Figure 1A). We therefore designated this structure the early cell mass (ECM). However, we do not know how the early cell mass develops an organised prothallus nor do we know how the prothallus initiates formation of the first meristem (Figure 1A).

**Figure 1:**
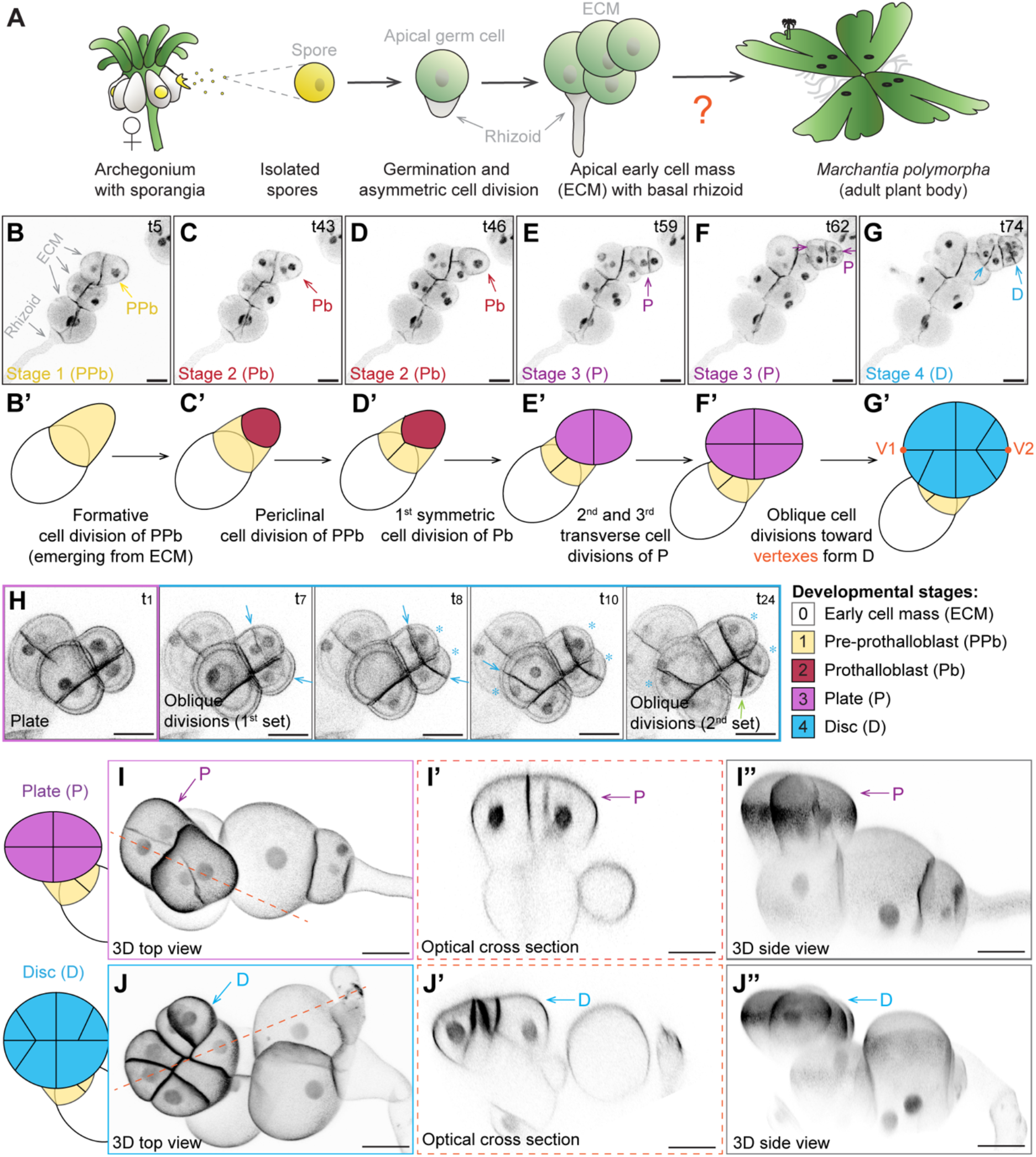
Reproducibly oriented cell division patterns initiate the 3D Marchantia plant body. Schematics on Marchantia development from spores: sporangia develop on archegonia and release single-celled spores that develop into sporelings. **B-G**) Confocal time lapse imaging of a 4 day-old sporelings (n=14) expressing the nuclear and plasma membrane fluorescent marker *pro*Mp*ROP:mScarlet-NLS*; *pro*Mp*UBE2:mScarlet-AtLTI6* ^14^. The sporeling was imaged for 88 h with 1 h frames. Selected time points (t5-t74) are depicted to highlight key developmental stages 1-4. From left to right: formation of the pre-prothalloblast (PPb, yellow arrow, t5, stage 1), formative division of the PPb forms an outer prothalloblast daughter that elongates while the inner daughter cell divides periclinally (Pb, red arrow, t43-t46, stage 2), three symmetric divisions of the Pb, two perpendicular to the first (transverse), form a 4-celled plate (P, purple arrow, t59-t62, stage 3), aligned oblique divisions initiate the disc (blue arrow, t74). B’-G’ show schematic and colour-coded representations of the early sporeling stages 1-4 with terminology of emerging cell identities. Scale bars 20 μm. The images shown represent time points from video S1. **H**) Confocal time lapse imaging of a sporeling between day 4-5. Sporelings were imaged for 24 h with 30 min frames. Oblique divisions (blue arrows) initiated formation of the disc. Scale bars 20 μm. Shown are time points taken from videos S2 and S3. **I-J**) 3D modelling of 4 day-old (plate stage, I-I”) and 5 day-old (disc stage, J-J”) sporelings expressing *pro*Mp*ROP:mScarlet-NLS*; *pro*Mp*UBE2:mScarlet-AtLTI6* using MorphographX ^36^. Top views of plate and disc are shown in I-J, side views of optical cross sections along the dotted red line are shown in I’-J’, side views of the same 3D reconstructed sporelings are shown in I”-J”. Scale bars 20 μm. Rotations of these 3D reconstructions are shown in videos S4 and S5.

Here we describe the developmental transitions that occur as the early cell mass develops a flat prothallus organ on which the first meristem initiates. Using 4-D imaging, 3-D prothallus reconstructions, patterns of gene expression and mutants, we define all the stages of sporeling development from the unicellular spore to the prothallus and the first meristem at cellular resolution. We demonstrate that the Marchantia Class III Homeodomain-Leucine-Zipper (MpC3HDZ) transcription factor marks the dorsal side of these initial flat body structures and establishes dorsoventrality before formation of the first meristem. These data describe the cell division patterns and molecular processes that generate a dorsoventral prothallus, on which the first meristem with an apical stem cell develops de novo.

## RESULTS

### Reproducibly oriented cell division patterns initiate the 3D Marchantia plant body

An asymmetric cell division of the spore cell produces a basal cell that differentiates as a primary rhizoid cell, and an apical germ cell that divides to form a morphologically heterogenous early cell mass (ECM) ^7,14^. To define the cell division pattens that form the flat prothallus body, we tracked cell divisions of 3 days-old sporelings expressing the nucleus and plasma membrane (PM) markers *pro*Mp*ROP:mScarlet-NLS*; *pro*Mp*UBE2:mScarlet-AtLTI6*, respectively ^14^ using confocal time lapse imaging. The most apical cell of the early cell mass, opposite the primary rhizoid cell (Figure 1A-B), elongated and formed a protrusion we designated as pre-prothalloblast as it initiated the prothallus (Figure 1B). The pre-prothalloblast underwent a formative division that generated a roundish cell towards the outside and a narrow cell on the inside (Figure 1C). The narrow daughter cell expanded and divided periclinally, forming a reproducible V-shaped cell wall junction (Figure 1D). The prothallus developed from the round outer cell positioned distal to the V-junction, and we therefore designated it the prothalloblast (Figure 1B). The prothalloblast divided once symmetrically along its shortest axis to form two cells (Figure 1E). Each of these cells divided once more transversely to generate four equally sized cells that together constituted a plate (Figure 1F). Three out of the four plate cells divided obliquely to generate smaller daughter cells that expanded the plate into a disc (Figure 1G and video S1). These data demonstrate that a single cell in the early cell mass gives rise to the prothallus.

The plate and disc are the first flat 3D prothallus structures that form in the haploid phase of the Marchantia life cycle. They can be approximated with an ellipsoid with a major axis connecting the two vertices (Figure 1F’-G’ and video S2). As the plate expanded into the disc, the first set of oblique divisions that formed the disc were temporally synchronised and aligned at the major axis (marked by blue arrows in Figure 1H). These oblique divisions generated smaller daughter cells that flanked the vertices of the disc (see blue asterisks in Figure 1H). The larger daughter cells divided without expanding during interphase and formed increasingly smaller cells at the expanding margin of the disc (green arrow in Figure 1H and video S3). To determine the 3D organisation of the sporeling plate and disc stages, 3D reconstructions of z-stacks were generated. Sporeling plates (Figure 2I-I” and video S4) and discs (Figure 2J-J” and video S5) formed spatially separated from the early cell mass. These data demonstrate that a pre-prothalloblast develops through anisotomous expansion of a cell in the early cell mass that undergoes a formative division to generate the prothalloblast. A flat prothallus forms through a reproducible pattern of transverse and oblique cell divisions of the prothalloblast.

**Figure 2:**
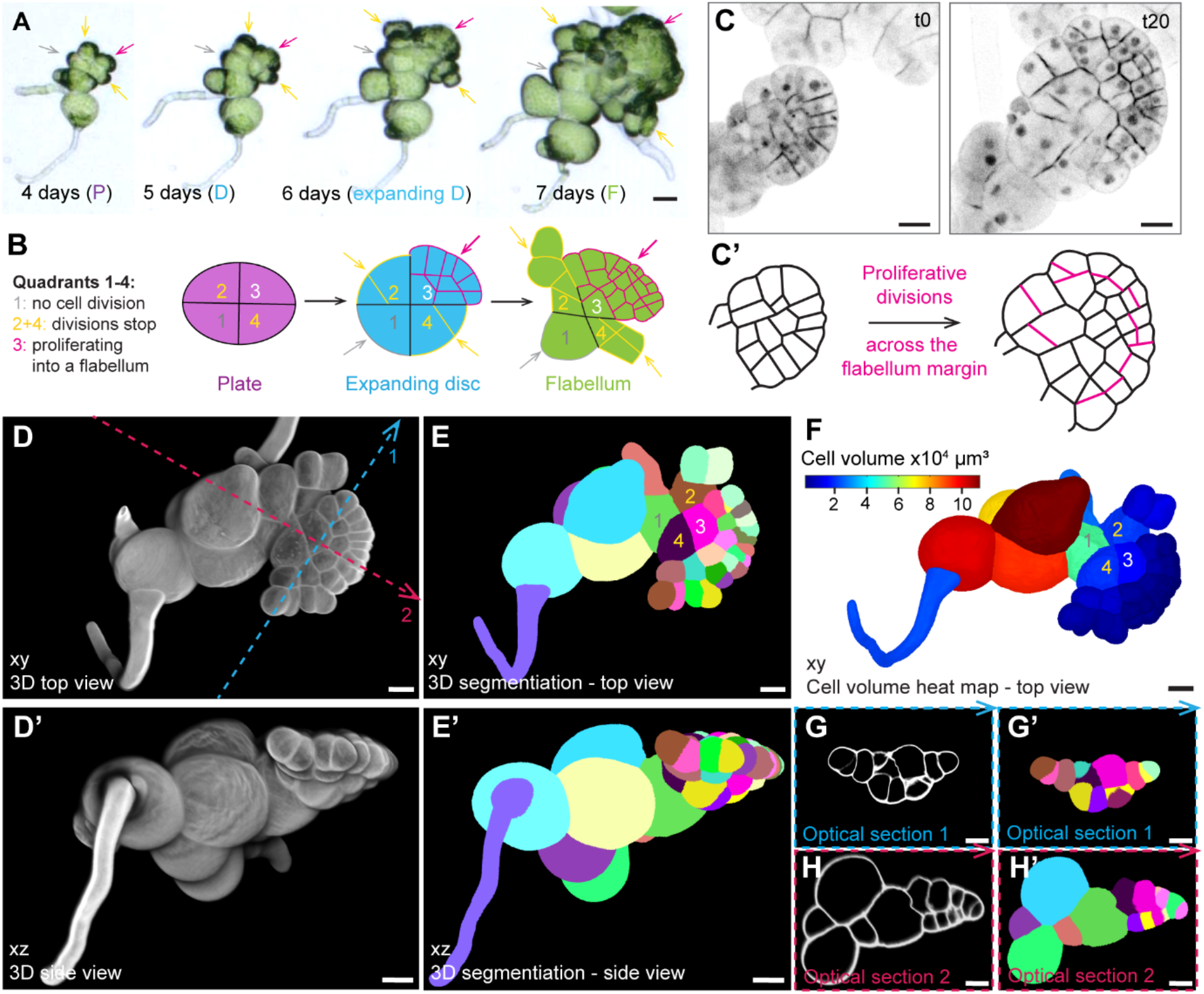
Flabella originate from single plate quadrants and proliferate into saucer-like 3D structures. **A)**Time-course imaging of flabellum development originating from plate quadrants (arrows at 4 days): one quadrant was dormant (grey arrow), two quadrants divided 1-3 times to expand into a disc while the most apical quadrant proliferated into a flabellum. N=40, scale bars 50 μm. **B)** Schematic representation of the disc to flabellum transition with proliferative potential marked by coloured arrows as described in F. **C**) Confocal time lapse imaging of a sporeling between day 8 to 9 expressing the nuclear and plasma membrane marker *pro*Mp*ROP:mScarlet-NLS; pro*Mp*UBE2:mScarlet-AtLTI6*. Sporelings were imaged for 20 h with 1 h frames. Schematic representations of the cell outlines are depicted in C’ and show that all cells along the flabellum margin divide (n=12). The depicted time points are based on video S6. Scale bars 20 μm. **D-D’**) 6 day-old sporelings (n=10) were cleared and cell walls stained with SR2200. A 3D reconstructed sporeling is shown from the top (xy, in D) and the side (xz, in D’). Blue and pink dotted lines indicated sites of optical cross sections as shown in G-H. Scale bars 20 μm. **E-E’**) Depicted are the sporelings shown in D after segmentation with MorphographX. Each coloured patch represents an individual cell of the sporeling. Scale bars 20 μm. **F**) A cell volume heat map of the sporeling that is shown in D and E. Rhizoid and all cells of the fan were coloured in blue, which represents a small cell volume. Scale bars 20 μm. **G-H’**) Shown are optical cross sections through the flabellum as indicated by dotted lines in D. Transverse sections (blue dotted line 1) show that the flabellum was 2-3 cells thick in the centre with a unicellular margin (G-G’). Sagittal sections (pink dotted line 2) show that the flabellum was attached to a large early cell mass cell (H-H’). The directions of the arrowheads indicate the orientation of the optical cross sections. G-H present optical sections with cell wall staining as shown in D, while G’-H’ depict the same optical sections after segmentation as shown in E. Scale bars 20 μm.

### Flabella originate from single plate quadrants and proliferate into saucer-like 3D structures

The sporeling plate generates a disc that in turn develops into a fan-shaped structure that we designate the flabellum. Since plant cells are immobile and encased by cell walls, we hypothesised that the cell lineages that form the semi-circular flabellum may only originate from a subset of the four plate cells. To determine the contribution of each plate cell to the flabellum, we tracked the development of plates over time (Figure 2A) .Typically, three of the four plate-cells – usually those furthest away from the primary rhizoid – divided obliquely to contribute to the disc (Figure 2A-B). The descendants of one plate cell formed most of the cells in the flabellum (pink arrow in Figure 2A-B). The two flanking plate-cells divided but cell divisions ceased in their descendants (yellow arrows in Figure 2A-B). The fourth plate cell rarely divided but expanded and did not contribute to the flabellum (grey arrow in Figure 2A-B). Therefore, each of the four plate cells constitute a quadrant with distinct proliferative potential (see schematic in Figure 2B). As depicted in our schematic, quadrant 1 did not contribute to the flabellum, while the opposite positioned quadrant 3 was highly proliferative and produced most of the expanding flabellum (Figure 2B). The current literature suggests that the first apical stem cell with two-cutting faces develops within this highly proliferative plate quadrant ^6,8,15^. An alternative hypothesis is that the flabellum first expands before a notch with meristem develops ^10^. To differentiate between these two hypotheses, we imaged developing flabella and traced all newly emerging cells for 20 hours using time lapse imaging (video S6). Cells across more than half of the flabella margin continued to divide during this period (Figure 2C-C’). These cell divisions contribute to new margin cells and cells inside the margin. No localised regions of cell division – characteristic of an apical cell and associated meristem – were observed. These data are consistent with the hypothesis that the increase in size of the flabellum is the result of cell proliferation along its margin as proposed by O’Hanlon in 1926 ^10^ and that the flabellum is not derived from a single apical stem cell or meristem, as postulated by Leitgeb 1881 ^4^ and currently held and cited ^8^.

To define the spatial relationships between the different regions of the sporeling, we reconstructed cell wall-stained, 6 day-old plants in three dimensions. The flabellum was flat and saucer shaped and protruded from the early cell mass (Figure 2D-D’). Cell volumes (Figure 2F) were smaller (≤ 2×10^4^ μm³) throughout the flabellum (blue colour in Figure 2F) than in the early cell mass (≥ 6×10^4^ μm³). All small flabellum cells could be traced back to three of the four plate quadrants, with the third contributing most cells, which was consistent with the observations in Figure 2A-B. Transverse optical cross sections that run perpendicular to the bilateral symmetry (apical-basal) axis of the flabellum (blue dotted line 1 in Figure 3D) indicated that the central region of the flabellum was two cell layers thick, while a single layer constituted the outer margin (Figure 2G-G’). The flabellum is two cell layers thick when viewed in the sagittal plane and was attached to the enlarged and undivided first plate quadrant at its base (green cell in Figure E, E’ and H’). These data support the hypothesis that the flabellum extends by marginal cell division and is not derived from a meristem.

**Figure 3:**
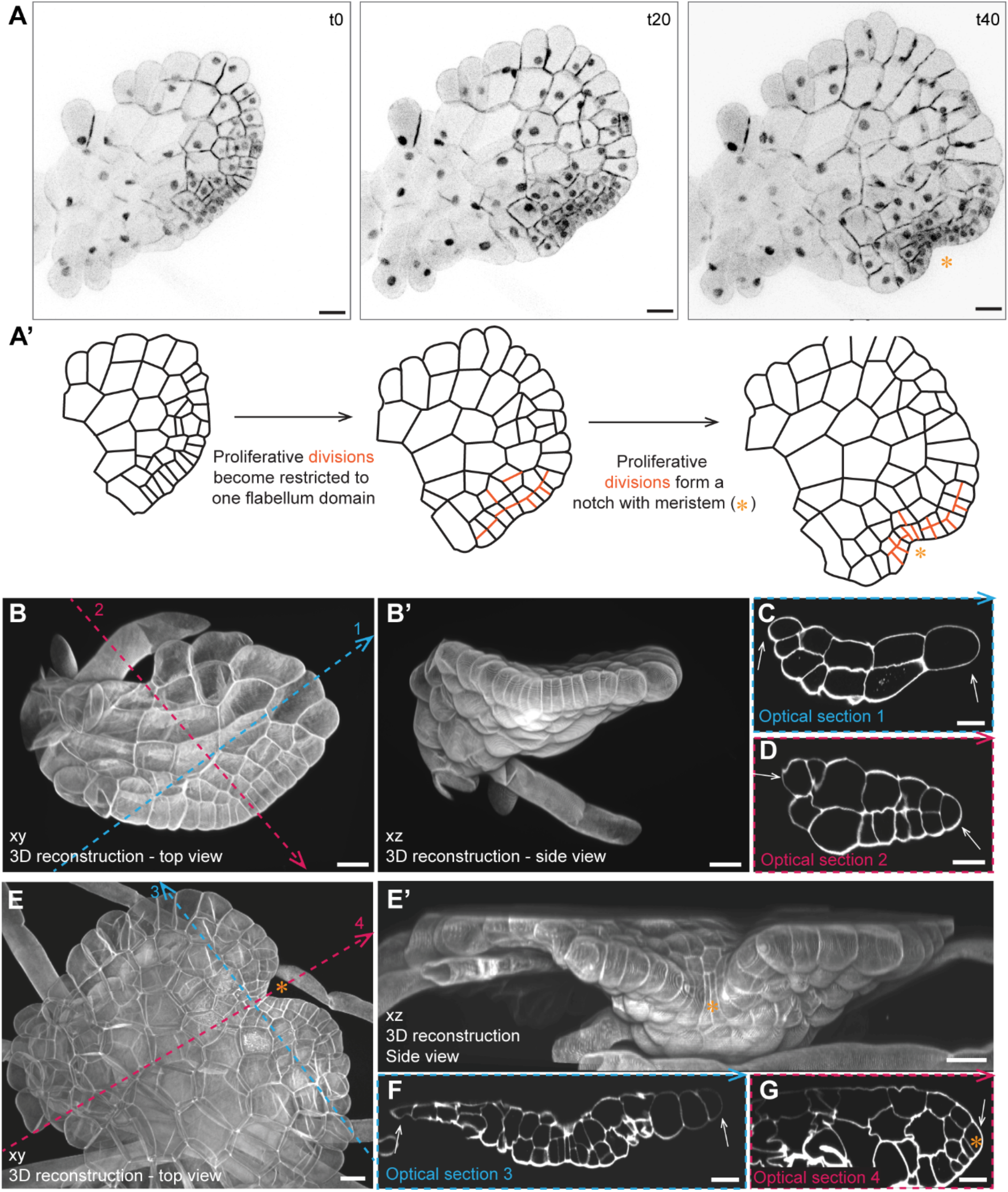
Localised cell division in the expanded flabellum predict the position of the first meristem. **A)**Confocal time lapse imaging of a sporeling between day 10 to 13 expressing the nuclear and plasma membrane marker *pro*Mp*ROP:mScarlet-NLS; pro*Mp*UBE2:mScarlet-AtLTI6*. Sporelings (n=6) were imaged for 59 h with 1 h frames and a 5 h gap at time point t38. Schematic representations of the cell outlines following the plasma membrane marker in the confocal images are depicted in A’. Within the expanded flabellum (t0), cell divisions became restricted to one domain of the flabellum (t20, orange lines), which eventually formed a notch characteristic for a Marchantia meristem (t40, orange asterisk). Depicted time points are based on video S7. Scale bars 50 μm. **B-G**) 10 day-old sporelings (n=7) were cleared and cell walls stained with SR2200. 3D reconstructed flabella without notch (B-D) and with notch (E-G) are depicted from the top (B, E) and the side (B’, E’). Dotted lines indicate sites of transverse (blue, C, F) and sagittal (pink, D, G) optical sections. Flabella without notch are 2 cell-layers thick with a unicellular margin (white asterisk, C-D) while flabella are 3-4 cell layers thick once a notch with wedge-shaped apical stem cell is present (orange asterisk in G). The direction of the arrowheads indicates the orientation of the optical cross sections. Scale bars 20 μm.

### Localised cell division in the expanded flabellum predict the position of the first meristem

The meristem of the mature plant is located in a notch that develops between two lobes of thallus tissue ^7^. To determine how and when the first notch forms, we tracked notch development by time lapse imaging of fully expanded flabella (Figure 3A). Expanded flabella formed a semi-circular, convex margin without a notch if viewed from the dorsal side (t0 in Figure 3A-A’). Along that convex margin, cell divisions became locally restricted and produced a niche of cells that were smaller than the surrounding cells (t20 in Figure 3A-A’). This highly proliferative niche encompassed two to three small cells flanked by several margin-cells to their left and right that divided periclinally and expanded faster than the small cells in the centre (t40 in Figure 3A-A’). This differential cell division and expansion pattern within the niche generated first a flattened margin (t20 in Figure 3A-A’) and later a concave indentation, that in due course developed into a notch (t40 in Figure 3A-A’). Consequently, notch formation involved the development of a localised region of cell division forming a population of small cells surrounded by tissue comprising relatively larger cells, as predicted by ^10^. Therefore, the formation of the meristem and the notch were not derived from a single apical stem cell as is the current view ^8^.

Flabella are two cell layers thick when they first develop (Figure 2D-H). However, the mature thallus is a multilayered tissue that is proposed to be generated from an apical stem cell with four cutting faces ^7,8^. To investigate if a notch indicated presence of such an apical stem cell and if its formation led to tissue thickening, we reconstructed expanded flabella in 3-D before and after notch formation (Figure 3B-G). Fully expanded, “round” flabella were flat and at least two cell layers thick with a one-cell thick margin (Figure 3C-D). After notch formation, the number of cell layers increased towards the future ventral side of the flabellum, which could be consistently located by rhizoid positioning (Figure 3E-E’). A transverse optical cross section showed that there were more cell layers around the notch than in the flabellum margins on either side, which remained one-cell thick (white arrows in Figure 3F). A sagittal section along the bilateral symmetry axis and through the centre of the notch showed that tissues around the notch were three to four cell layers thick (Figure 3G). A wedge-shaped epidermal cell, whose position and morphology are consistent with that of an apical stem cell in thallus meristems ^7^, separated the dorsal (upper) and ventral (bottom) sides of the prothallus (Figure 3G). These data demonstrate that the first meristem with apical stem cell forms de novo in notches generated by spatially restricted proliferation and cell expansion in one region of the flabellum and that tissue thickening towards the ventral side correlates with emergence of a notch with an apical stem cell.

### Mp*C3HDZ* activity is required for the development of a flat and dorsoventralised Marchantia thallus

Having discovered that the meristem forms on a flat but multilayered flabellum, we hypothesized that the molecular mechanism that establishes dorsoventrality and prothallus flatness could be already active before meristem initiation. As a first step, we investigated if dorsoventrality regulators are conserved between *Arabidopsis thaliana* and *Marchantia polymorpha*. The Class III HD-Zip (C3HDZ) transcription factor family promotes dorsoventrality of flat angiosperm organs, such as leaves ^16^ and had been associated with phylid development in the moss *Physcomitrium patens* ^17^. There is a single *C3HDZ* gene orthologue (Mp1g24140) in Marchantia. We reasoned that if this transcription factor was functionally conserved across land plants, loss of function mutants of MpC3HDZ should be defective in developing dorsoventrality and flat thalli. To test this hypothesis, we generated three loss-of-function mutants using CRISPR-mediated gene editing, Mp*c3hdz-1*, Mp*c3hdz-2* and Mp*c3hdz-3* (Figure 4A). In Mp*c3hdz-1* there was an in-frame 12 bp deletion at positions 158-170 bp that resulted in the deletion of the four amino acids Leu, Ala, Asn and Ile within the homeobox domain In Mp*c3hdz-2* there was a 2 bp deletion at position 159-160 bp that caused a frame shift in the homeodomain coding sequence and a premature stop codon at position 300 bp. Mp*c3hdz-3* was characterised by a 7 bp deletion at position 362-368 bp, which led to a frame shift and a premature stop codon at position 375 bp (Figure 4A). All three Mp*c3hdz* mutants developed similar phenotypes of radialised, stalk-like thalli (Figure 4B-C). Wild type plants developed flat, thin thalli with a clear separation between dorsal and ventral sides (Figure 4B-D). By contrast, Mp*c3hdz* developed almost cylindrical, largely radially symmetric axes (Mp*c3hdz-1* and Mp*c3hdz-2*) or occasionally fused (Mp*c3hdz-3*) thalli margins with disorganised tissue outgrowths on their surfaces (Figure 4D). These data indicate that MpC3HDZ regulates tissue flatness in Marchantia.

**Figure 4:**
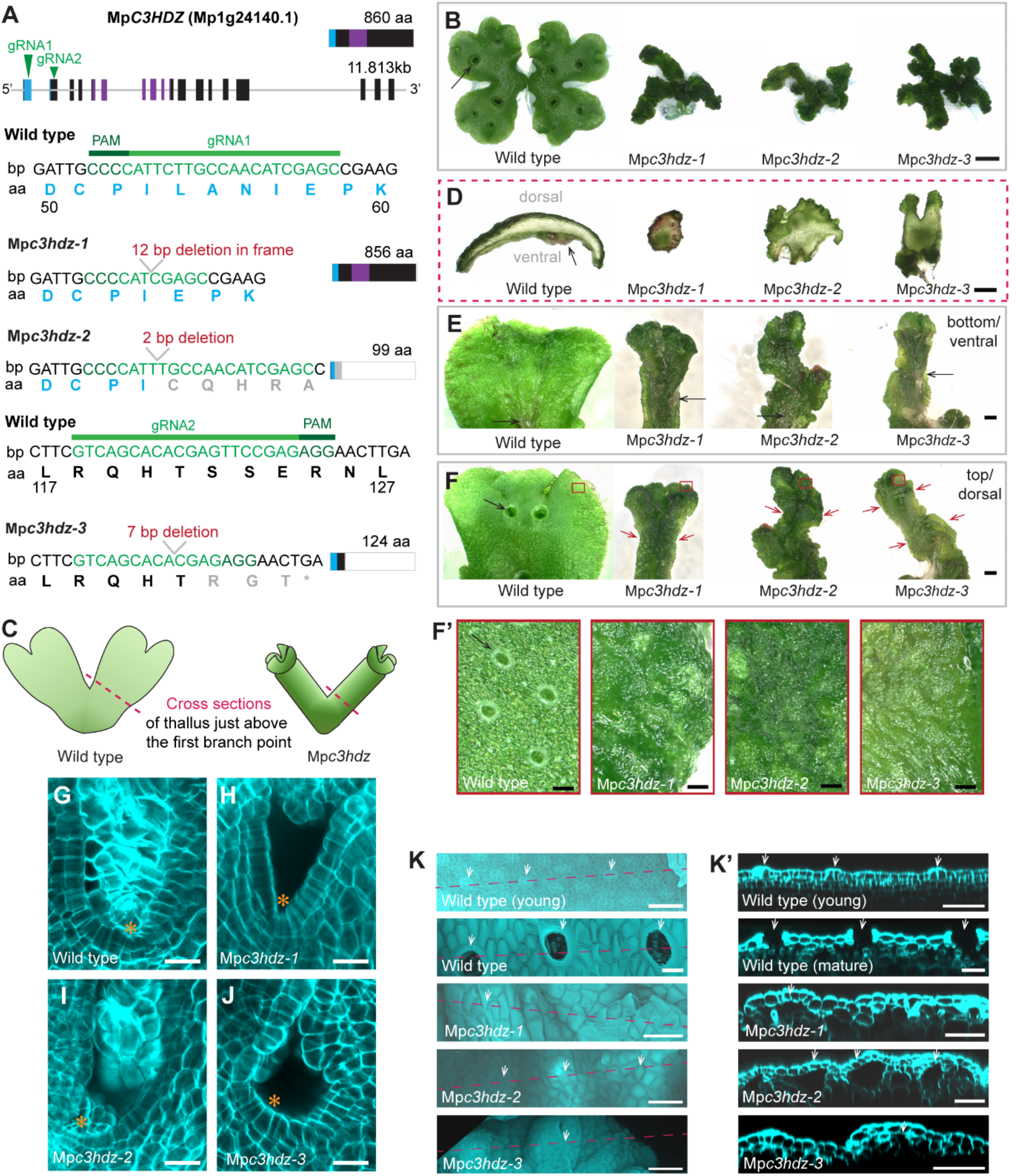
Mp*C3HDZ* activity is required for the development of a flat and dorsoventralised Marchantia thallus. **A)**Schematic representation of the longest *C3HDZ* splice variant transcript Mp1g24140.1 (11.813kb) and the corresponding protein (860 aa) with the homeobox domain indicated in blue and the START (StAR-related lipid-transfer domain) coloured in purple. Two gRNAs were designed. gRNA1 targeted the homeobox domain and gRNA2 a region after the homeobox but before the START domain. Base pair and translated amino acid sequences are depicted for each gRNA binding site in wild type and the selected mutants: Mp*c3hdz-1* has an in-frame 12 bp deletion in the homeobox domain, Mp*c3hdz-2* has a 2 bp deletion in the homeobox domain resulting in a premature stop codon and Mp*c3hdz-3* has a 7 bp deletion after the homeobox domain resulting in a premature stop and truncated protein. **B)** 16 day-old Tak1xTak2 plants grown from a gemma alongside Mp*c3hdz-1*, Mp*c3hdz-2* and Mp*c3hdz-3* mutants grown from thallus stubs as all Mp*c3hdz* mutants were infertile and produced no gemma cups (black arrow). Scale bar 3 mm. **C-D)** Hand cross sections were taken from 4 week-old thallus tissue close above the first branch point as indicated by a dashed pink line in the schematic (C). Cross sections of wild type thallus were thin and broad with a separated dorsal (air pores) and ventral (mid rib and rhizoids, black arrow) side, while cross sections of Mp*c3hdz* mutants were roundish with radialised tissues and no obvious tissue types that could indicate dorsoventrality. Scale bars 1 mm. **E-F)** Bottom (ventral, E) and top/dorsal (F) thalli of 4 week-old wild type (Tak1xTak2) and Mp*c3hdz-1*, Mp*c3hdz-2* and Mp*c3hdz-3* mutants. Close-up images (red squares) of dorsal tissues at the thallus apex show a flat epidermis with gemma cups (arrow in E) and air pores in wild type but bumpy epidermises without gemma cups or apparent air pores in Mp*c3hdz* mutants (F-F’). Scale bars 1 mm (E) and 100 μm (F). **G-J)** Optical cross sections of cleared and cell wall-stained Mp*c3hdz* mutants. All Mp*c3hdz* allels possessed notches with meristems that produce small cells (orange asterisks in H-J) at the apices of their trumped-shaped thalli. These meristems did not differ morphologically from wild type (G). Scale bars 20 μm. **K)** Depicted are cross sections of dorsal epidermis. Air pores, a typical dorsal structure, were closed in the young wild type epidermis close to the meristem and open in the mature epidermis (white arrows, see also F-F’). All Mp*c3hdz* mutant alleles were missing obvious air pores but possessed air chambers underneath convex epidermal bumps in trumpet-like folds at their thallus apices (white arrows in K’). Scale bars 50 μm.

In angiosperms, C3HDZ family members regulate meristem formation alongside abaxial-adaxial identity^16,18^. We tested the hypothesis that meristem formation was defective in Mp*c3hdz-1*, Mp*c3hdz-2* and Mp*c3hdz-3* mutants. Active meristems were visible as domains of smaller, dividing cells within the notches of all cleared and wall-stained Mp*c3hdz* mutants (orange asterisks in Figure 4G-J). Taken together these data indicate that C3HDZ protein is required for the development of dorsoventrality but not for meristem initiation in Marchantia.

To investigate if dorsal and ventral tissue identities could still be detected in Mp*c3hdz*, we defined the top (dorsal) and bottom (ventral) sides of the radialised Mp*c3hdz* mutant thalli relative to their growth orientation on culture plates and examined each side in detail. Just like wild type, all Mp*c3hdz* mutants developed midrib rhizoids on their ventral sides (Figure 4E). However, the dorsal side of Mp*c3hdz* thalli was fused into a trumpet-like shape and typical dorsal structures, such as gemma cups and air pores were absent (Figure 4F-F’). This suggests that MpC3HDZ protein activity is required for the development of dorsal tissue identities. Although there were no air pores within the Mp*c3hdz* apices in the trumpet-shaped regions (red squares in Figure 4F-F’), we hypothesised that the inner tissues of the trumpet-shape Mp*c3hdz* apex may develop dorsal-like identity. Each Mp*c3hdz* mutant developed an uneven dorsal surface – comprising convex surface protrusions and concave hollows that we refer to as bumps – while wild type dorsal surface is flat with air pores (white arrows in Figure 4K). The young wild type epidermis close to the meristem developed air pore complexes with initially closed apertures that later opened. An air chamber developed underneath these complexes in mature tissue (white arrows in Figure 4K-K’). Optical cross sections through Mp*c3hdz-1*, Mp*c3hdz-2* and Mp*c3hdz-3* mutant dorsal regions showed that air chambers were present underneath convex bumps of the epidermis, but air chamber roofs did not form pores (white arrows in Figure 4K’). Dorsal tissue development is therefore defective in these mutants. Based on these data we conclude that Mp*c3hdz* mutants are largely ventralised in the cylindrical regions but develop small regions of defective dorsal identity at the apices of the trumpet-shaped folds. These data demonstrate that Mp*C3HDZ* controls the development of dorsal tissue formation, organ flatness and dorsoventrality in Marchantia thalli.

### The MpC3HDZ transcription factor accumulates in cells of the first flat body structures

To determine if dorsoventrality was established in the prothallus before meristem initiation, we imaged sporelings expressing *pro*Mp*C3HDZ:*Mp*C3HDZ-VENUS* together with the PM and nuclear marker *pro*Mp*ROP:mScarlet-NLS; pro*Mp*UBE2:mScarlet-AtLTI6*. In 3 day-old sporelings, MpC3HDZ-VENUS fluorescence was occasionally observed in the nucleus of the primary rhizoid or weakly in older early cell mass cells, but never in the prothalloblast (Figure 5A-A’). There was weak VENUS fluorescence in the four-celled plate (Figure 5B-B’) and stronger VENUS fluorescence in the nuclei of disc cells (Figure 5C-C’). This indicated that MpC3HDZ-VENUS signals increased in intensity at the plate to disc transition (Figure 5B-C and video S9). MpC3HDZ-VENUS fluorescence was brightest in the disc quadrant that formed the expanding flabellum (Figure 5D-D’ and Figure 3A) and remained strong in the flabellum (Figure 5E-E’) until signals became restricted to the proliferation zone around the notch as the meristem was initiated (Figure 5F-F’). We hypothesised that the expression of MpC3HDZ-VENUS would be polarized within the flabellum, the first dorsoventralised, flat structure comprising more than one cell layer (Figure 3). To test this hypothesis, we took optical cross sections through cleared flabella expressing *pro*Mp*C3HDZ:*Mp*C3HDZ-VENUS*. MpC3HDZ-VENUS fluorescence was greatest in the dorsal cell layer in multilayered regions of the flabellum. MpC3HDZ-VENUS fluorescence was also strong at the one-cell thick flabellum margin but never observed in the ventral or internal tissues of the multilayered flabellum (Figure 5G). Later, after the formation of a notch with an active meristem (white arrows in Figure 5H), MpC3HDZ-VENUS fluorescence remained polarised in cell layers of the dorsal tissues (white arrows in Figure 5H). We conclude that MpC3HDZ-VENUS marks flat structures throughout prothallus development and the dorsal sides of flabella and through to the stage where a de novo meristem is active in the notch.

**Figure 5:**
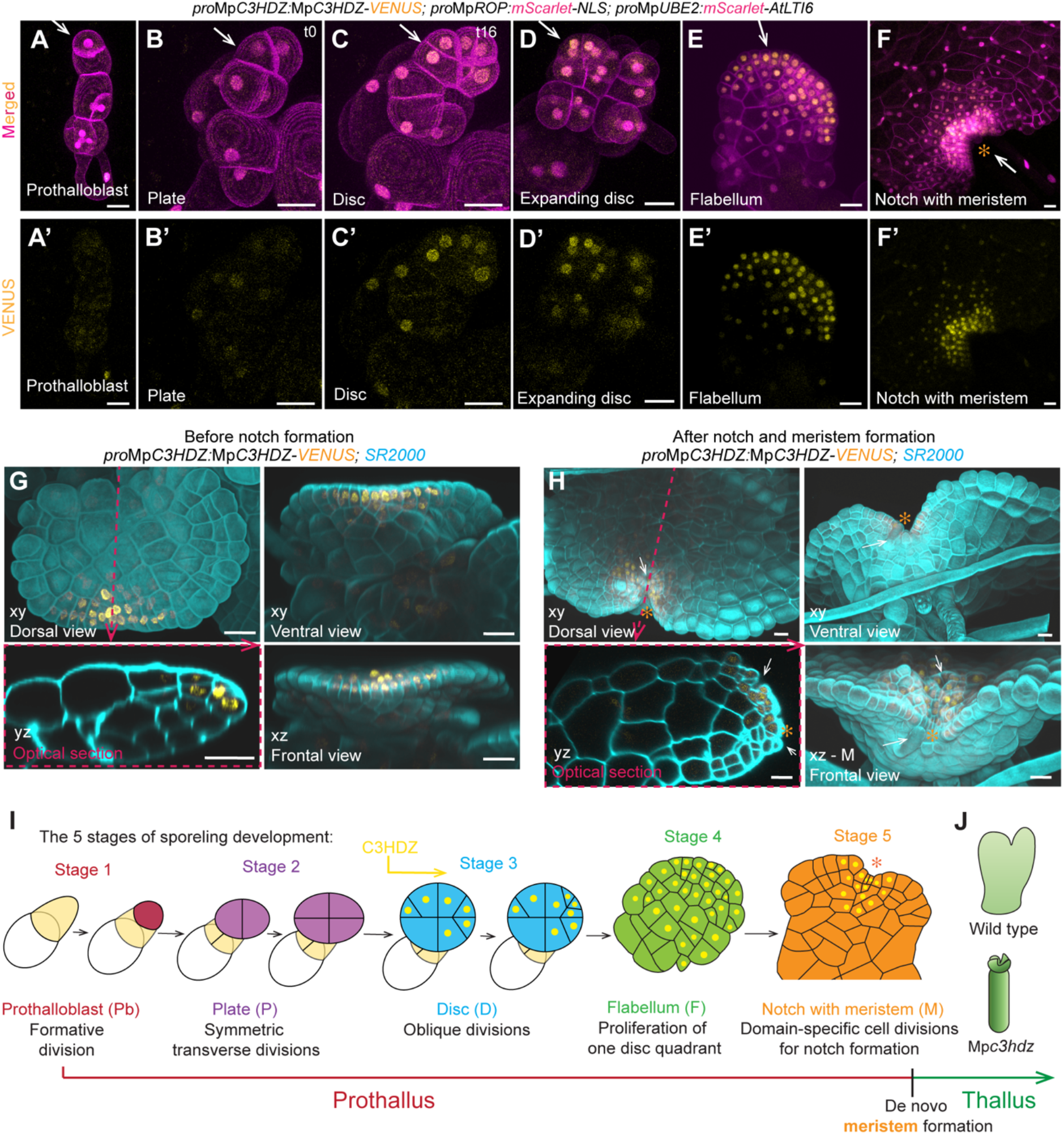
The MpC3HDZ transcription factor accumulates in cells of the first flat body structures. **A-F)** Confocal images of sporelings expressing *pro*Mp*C3HDZ:*Mp*C3HDZ-VENUS* with plasma membrane and nuclear marker *pro*Mp*ROP:mScarlet-NLS; pro*Mp*UBE2:mScarlet-AtLTI6* at key developmental stages of the prothalloblast (n=12 in A-A’), the plate (n=5 in B-B’) and its transition during 16 h time lapse imaging into a disc (n=4 in C-C’, see video S8), the proliferating disc quadrant (n=15 in D-D’), the expanded flabellum (n= 32 in E-E’) and the meristem (n=7 in F-F’). Clear C3HDZ-VENUS signals were first observed in the disc. Scale bars 20 μm. **G-H)** 8 to 9 day-old sporelings expressing *pro*Mp*C3HDZ:*Mp*C3HDZ-VENUS* were cleared and cell walls stained with SR2200. Depicted is a 3D reconstructed expanded flabellum (n=7 in G) and a flabellum with the first meristem indicated by the notch (n= 10 in H) from the dorsal/top side (xy), the front (xz) and the ventral/bottom side (xy). Optical cross sections (yz, see dotted pink line) show MpC3HDZ-VENUS polarised to the dorsal side (G’-H’). The first differentiated tissues produced by the new meristem are indicated by white arrows (air chambers on the dorsal side and slime papillae on the ventral side in H). Scale bars 20 μm. **I)** Schematic representation of prothallus development with prothalloblast (stage 1), plate (stage 2), disc (stage 3), flabellum (stage 4), and de novo meristem (stage 5) that generates adult thalli. MpC3HDZ-VENUS first localises to nuclei after oblique divisions at the transition from plate to disc and signals intensify on the dorsal side of the expanding flabellum before being restricted to the dorsal side of the meristem. **J)** Mp*C3HDZ* promotes dorsoventrality and body flatness as Mp*c3hdz* mutants show thick radialised thalli that lost dorsoventrality and have defective dorsal tissue formation.

## Discussion

The haploid body of *Marchantia polymorpha* develops from a meristem that forms among a population of dividing cells – the early cell mass (ECM). We demonstrated that a single prothalloblast originates from the early cell mass to initiate development of a flat prothallus on which a meristem forms de novo. To generate the prothallus, the prothalloblast undergoes a regular pattern of formative, transverse and oblique cell divisions. We also discovered that the MpC3HDZ transcription factor regulates the development of dorsoventrality and polarises to the dorsal tissues of the prothallus before protein localisation becomes restricted to the dorsal side of the de novo formed meristem.

The prothalloblast sets a clear developmental boundary between the irregularly shaped early cell mass and the flat prothallus. Transverse divisions that form a quadrant of four cells had been described in early development of several liverworts, including *Blasia pusilla* ^6^, *Marchantia polymorpha* ^5,15^, *Fossombronia wondraczekii* ^19^, *Pressia quadrata* ^12^ and others ^20^. This suggests that forming a plate with four quadrants is a conserved developmental step in liverwort development. Tracking cell division orientation allowed us to further dissect this developmental stage: we discovered that the plate is product of transverse symmetric divisions derived from the prothalloblast, while the disc is a product of oblique divisions from a subset of plate quadrants. One quadrant of the disc proliferates into a flat prothallus structure we call the flabellum (Figure 5I). All cells of the flabellum undergo size-reducing periclinal and anticlinal divisions before divisions become restricted to a subset of marginal cells that form a notch through differential cell division and expansion as proposed by O’Hanlon ^10^. Alternatively, it had been hypothesised that the flabellum is product of an active apical stem cell with two cutting faces that generates a unistratous prothallus ^4,5,8,15^. Because of the resolution by which we could track cells and their descendants with modern imaging technologies, we are confident that this issue is now resolved.

Likewise, the establishment of dorsoventrality was so far based on the observation that an active meristem produces cell lineages that differentiate into morphologically distinct dorsal tissues (air pores and air chambers) or ventral tissues (rhizoids) ^8,10^. The expression of MpC3HDZ-VENUS suggests that dorsoventrality developed progressively from the single-cell-layered disc to the multi-layered flabellum. A wedge-shaped apical stem cell was first observed late in flabellum development after a notch was already formed. This indicates that the establishment of dorsoventrality can be observed on a molecular level long before the first morphologically distinct dorsal and ventral tissues are produced by the first meristem.

The Class III Homeodomain-Leucine-Zipper (C3HDZ) family is conserved among land plants and had been extensively studied in the angiosperm *Arabidopsis thaliana* ^16,18^. There are five *C3HDZ* homologues in Arabidopsis: *PHABULOSA* (*PHB*), *PHAVOLUTA* (*PHV*), *REVOLUTA* (*REV*), *ARABIDOPSIS THALIANA HOMEOBOX 8* (*ATHB8*) and *ATHB15*. Meristems do not develop in the triple *rev;phb;phv* mutants. These mutants also develop radialised ^16^ or – in weaker instances – radially fused, trumpet-shaped cotyledones and leaves ^18^. This trumpet-like phenotype is hypothesized to results from the development of a restricted dorsal (adaxial) domain on otherwise ventralised (abaxialised) mutant leaves ^18^. In Marchantia, the dorsal localisation of MpC3HDZ proteins seems to be essential for flat body formation because radialised and occasionally trumpet-shaped thalli developed in three independent Mp*c3hdz* mutants. However, all Mp*c3hdz* mutants developed meristems and could still branch, which suggests that Mp*C3HDZ* function is not required for meristem initiation or activity in *Marchantia polymorpha*. Consistent with our observations, the C3HDZ orthologue in the moss *Physcomitrium patens* regulates leaf (phylid) development but not meristem development ^17^. Together these independent observations support the hypothesis that C3HDZ protein function is required for the development of flattened dorsoventral structures in the haploid generation of bryophytes and the diploid generation of angiosperms.

## STAR METHODS

Detailed information about reagents and resources are listed in the Key Resource Table.

Plasmids and primer lists and light spectra are presented in Table S1.

## Key resources table

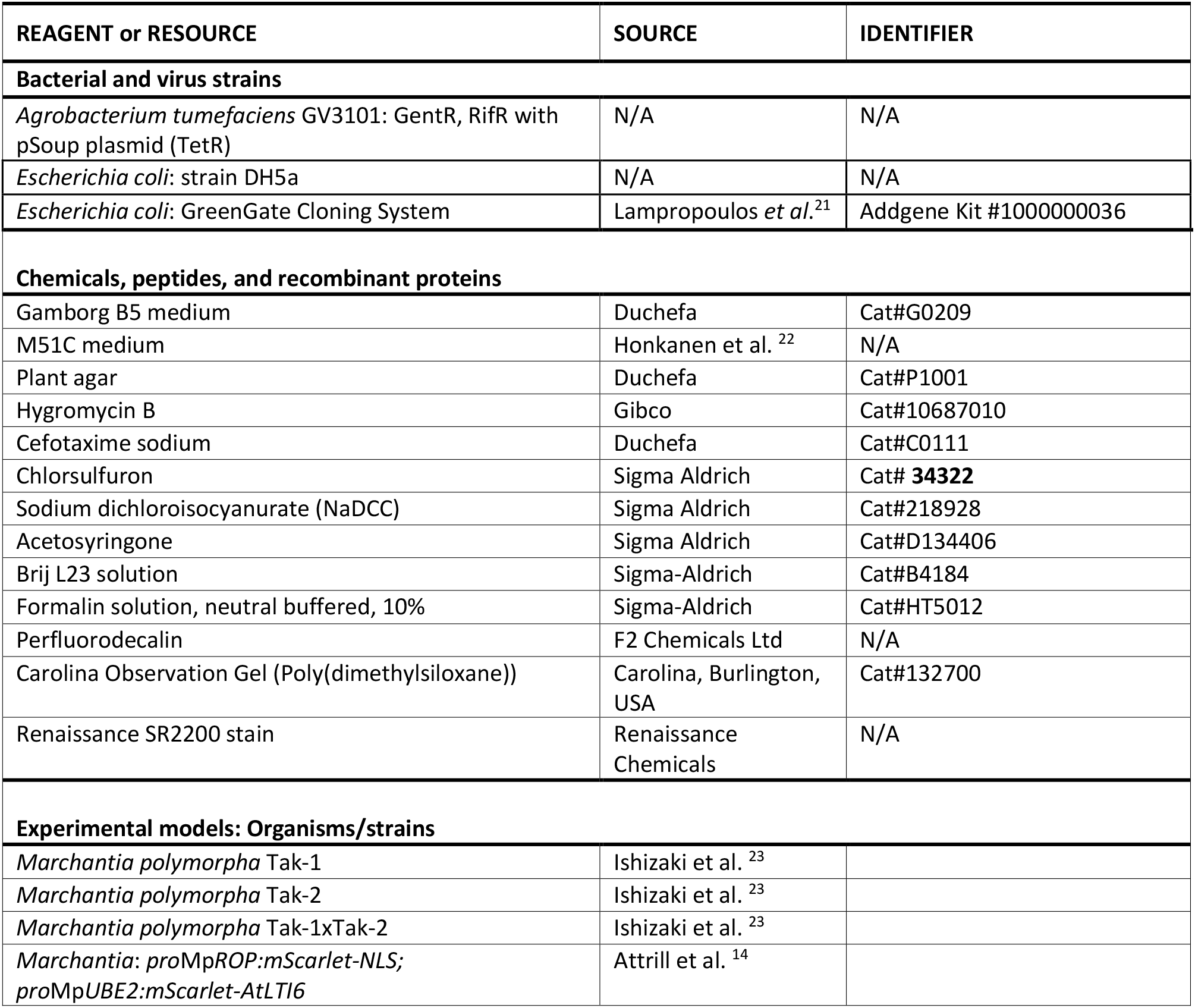

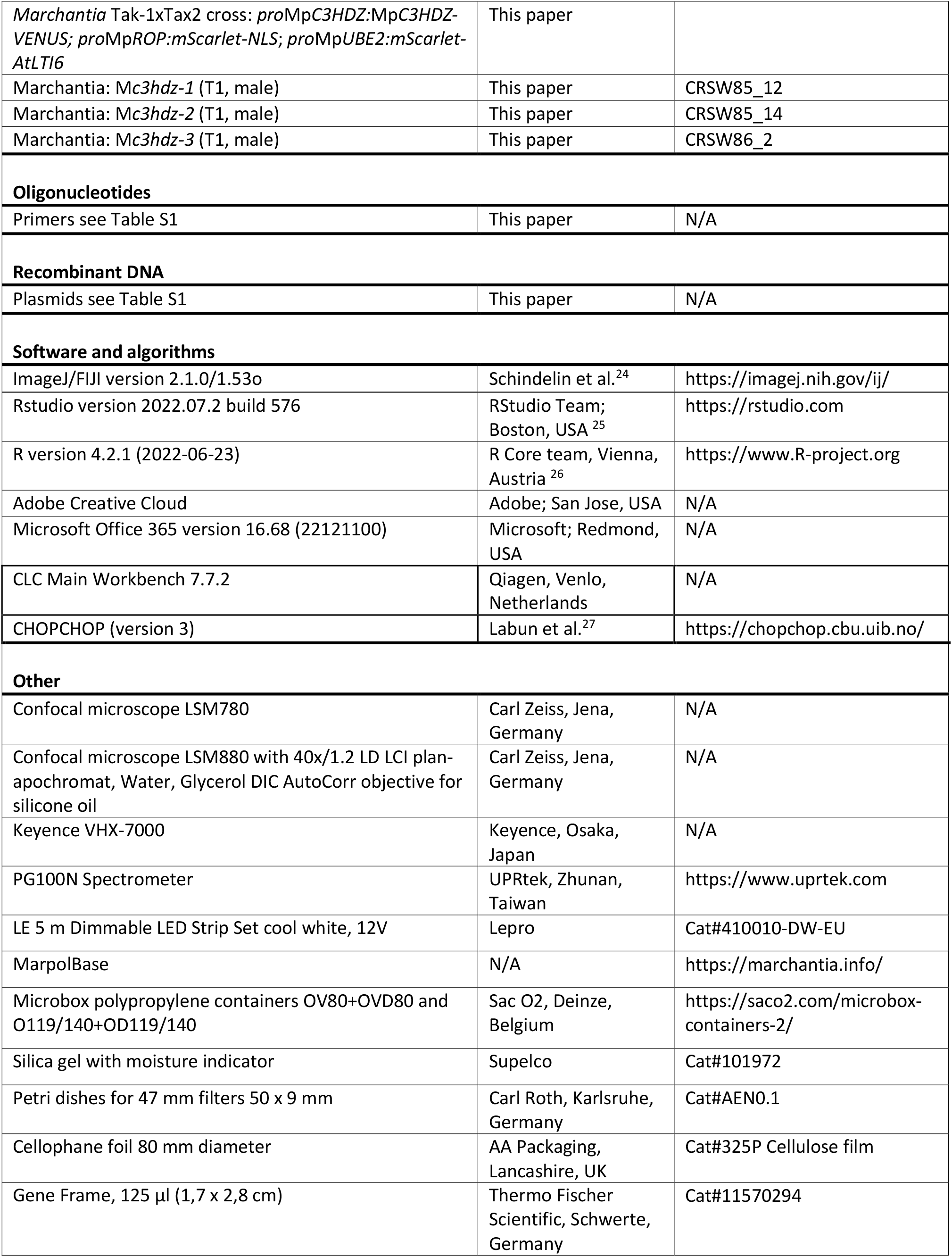

## Resource availability

### Lead contact

Requests for resources and further information should be directed to Liam Dolan.

### Materials availability

Newly generated lines and sources of previously reported transgenic lines are listed in the key resource table. For the transfer of transgenic material, appropriate information on import permits will be required from the receiver.

### Data and code availability

All data required to substantiate the claims of this paper are included in main or supplemental data. This paper does not report original code.

## EXPERIMENTAL MODEL AND STUDY PARTICIPANT DETAILS

*Marchantia polymorpha* wild type accessions Takaragaike-1 (Tak-1, male) and Takaragaike-2 (Tak-2, female) were used in this study ^23^. Plants from gemmae and thalli were grown in sterile petri dishes containing ½-strength B5 Gamborg’s agar (1.5 g/L B5 Gamborg, 0.5 g/L MES hydrate, 1 % sucrose, pH adjusted to 5.5 solidified with 1 % plant agar) at 23 °C under standard continuous white light (45 μmol m^-2^ s^-1^). To induce gametangiophore formation for crossing, 3 week-old gemmalings were transplanted into SacO2 Microbox containers with a autoclaved 1:3 mixture of vermiculite and Neuhaus N3 compost and grown under 50 μmol m^-2^ s^-1^ (PPFD) white light supplemented with 45 μmol m^-2^ s^-1^ far red light irradiation (spectra see Table S1) in a long day regime of 16 h light, 8 h dark at 23 °C ^28^.

## METHOD DETAILS

### Generation and germination of spores for sporeling analyses

Crosses between Tak-1 and Tak-2 were performed by releasing sperm from ripe Tak-1 antheridiophores in 200 μl of sterile water for 10 min before transferring the liquid onto immature Tak-2 archegoniophores ^28^. Based on ^23^ and as described in ^29^, mature but intact sporangia were harvested 3-4 weeks after crossing and dried in closed microbox polypropylene containers OV80+OVD80 filled with silicagel. After 4-5 weeks of drying, microcentrifuge tubes with single sporangia were directly frozen at -70°C for storage. To germinate spores, frozen sporangia were thawed and crushed in sterilization solution of 0.1% (w/v) sodium dichloroisocyanurate. After 2 min of sterilization, spores were pelleted by centrifugation at 13000 g for 3 min, the supernatant was removed, and spores resuspended in 500 μl sterile water. To grow sporelings for analysis and phenotyping, ½-strength B5 Gamborg’s agar (1.5 g/L B5 Gamborg, 0.5 g/L MES hydrate, 1 % sucrose, pH adjusted to 5.5 solidified with 1 % plant agar) was poured in small petri dishes (50 mm diameter, see key resource table) and topped with pre-cut, sterilised and water-wetted cellophane. Approximately 100 μl of spore suspension were plated per small petri dish, excess water was evaporated, petri dishes tightly closed and placed on black paper in a growth chamber with standard continuous white light conditions (45 μmol m^-2^ s^-1^). The black paper was critical to prevent light reflections and to ensure directional sporeling development towards a single light source.

### Generation of plasmids for plant transformation

Translational fusion construct *pro*Mp*C3HDZ:*Mp*C3HDZ-VENUS* was generated using the Green Gate method ^21^. Gene sequences from genomic DNA were amplified by Phusion PCR with Green Gate-specific primers listed in Table S1 and cloned via BsaI restriction sites into entry modules with a pUC19 based vector backbone. Final constructs (Table S1) with pGreen-IIS based vectors backbone were generated by assembly of entry modules according to the Green Gate method ^21^. The linker-VENUS module was published previously ^30^ and the chlorsulfuron plant resistance module was adapted from the OpenPlant toolkit ^31^.

To obtain mutant alleles Mp*c3hdz-1*, Mp*c3hdz-2* and Mpc3hdz*-3* by CRISPR/Cas9 mutagenesis, two gRNAs consisting of 20 nucleotides followed by a NGG sequence were designed to target the first and second exon of the Mp*C3HDZ* (**Mp1g24140/Mapoly0061s0107**) gene, respectively, using the ChopChop web tool ^27^. Constructs

SWCR85 (sgRNA 1) and SWCR86 (gRNA 2) were cloned by following instructions of the OpenPlant toolkit published by Sauret-Güeto et al.^31^

### Generation and origin of transgenic and mutant lines

Transgenes were introduced into wild type sporelings generated from a cross between Tak-1 and Tak-2 wild-type accessions of *Marchantia polymorpha*. Frozen sporangia were sterilised as described above and the suspension of one sporangium was cultured in sterile 125 ml Erlenmeyer flasks with 25 ml M51C medium ^23^ for 7 days at 23 °C under continuous white light (45 μmol m^-2^ s^-1^) with constant agitation (130 rpm). Electrocompetent Agrobacteria (see key resource table) were transformed with the desired plasmid and a dense overnight culture grown in 5 ml LB-medium with respective antibiotics from 2-3 colonies. The Agrobacteria culture was pelleted and the pellet resuspended in 10 ml M51C medium containing 100 mM acetosyringone and incubated for 6 h at 28°C. 1 ml of the Agrobacterium-induced M51C culture was added to the 7-day old sporeling culture and acetosyringone was added to a final concentration of 100 mM. Sporelings and Agrobacteria were co-cultivated for 2 days before the sporelings were washed with sterile water using a 40 mm nylon cell strainer and plated on ½-strength B5 Gamborg’s agar (1.5 g/L B5 Gamborg, 0.5 g/L MES hydrate, 1% sucrose, pH adjusted to 5.5 solidified with 1% plant agar) with 10 mg/ml cefotaxime and either 0.01 mM chlorsulfuron (*pro*Mp*C3HDZ:*Mp*C3HDZ-VENUS*) or 10 mg/ml hygromycin (CRISPR constructs) for selection. Plasma membrane and nuclear marker *pro*Mp*ROP:mScarlet-NLS; pro*Mp*UBE2:mScarlet-AtLTI6* (with plant hygromycin resistance) was generated and used before^14^ and served as “wild type” control for most confocal imaging experiments, unless stated otherwise. Crosses between *pro*Mp*C3HDZ:*Mp*C3HDZ-VENUS* and *pro*Mp*ROP:mScarlet-NLS; pro*Mp*UBE2:mScarlet-AtLTI6* were used for confocal imaging and time lapses.

### Imaging of living tissues

Wild-type sporelings and mutant plants were imaged with the Keyence VHX7000 digital microscope equipped with the VH-Z00R/Z00T and VH-ZST lenses and the VHX-7020 camera. For confocal imaging, cellophane pieces with sporelings growing on top were cut from the agar plate and freshly mounted with water in Gene Frame caskets followed by immediate imaging at an LSM780 (Zeiss) with 20x air or 40x water objectives. Lines expressing VENUS were excited by an argon laser at 514 nm and the emission was detected between 520-540 nm. mScarlet was excited at 561 nm (DPSS) and emission was collected between 600-650 nm.

### Time lapse imaging of sporelings

A self-made time lapse chamber was built based on instructions by Kirchhelle and Moore ^32^. To adapt this method for sporelings, 3 mm wide glass strips were cut from commercial objective slides with a glass cutter and four of these glass strips were glued with double sided tape onto a new objective slide to create a rectangular glass chamber. This chamber was sterilised with 70 % ethanol and filled with ½-strength B5 Gamborg’s agar (1.5 g/L B5 Gamborg, 0.5 g/L MES hydrate, 1 % sucrose, pH adjusted to 5.5 solidified with 1 % plant agar). A cover slip was placed temporarily on top during solidification of the agar to generate a smooth surface. To create extra space for the sporelings, a thin layer of gas-permeant poly(dimethylsiloxane) gel ^32^ was placed on top of the glass casket and any excess gum was used to seal the outer walls of the chamber. A small rectangle of cellophane (approx. 1.5 x 1.0 cm) with sporelings growing on top was cut from the agar plate, transferred on top of the solidified agar in the glass chamber and immediately topped with perfluorodecalin to prevent tissue from drying out. The glass casket time lapse chamber was sealed with a cover slip, gently pressed onto the poly(dimethylsiloxane) gel and secured with tape running around each side of the glass chamber. This time lapse chamber was taped onto the Zeiss LSM780 stage under an upright 20x/0.8 plan-apochromat air objective and cool white LED lights set to 100% power (45 μmol m^-2^ s^-1^, see Table S1) where attached around the stage. Sporelings were illuminated by the LED strip for 45 min with 15 min breaks in which z-stacks of 8-12 positions were taken, resulting in 20-90 h time lapses with typically 1 h frames (unless stated otherwise in the figure legends).

### Sporeling fixation, clearing, cell wall staining and imaging for 3D MorphographX reconstruction

The cellophane with sporelings (6-12 days-old) was peeled off the agar plate (see growth conditions for sporelings above) and immediately soaked in 1.5 ml fixative (10 % formalin solution with 0.1 % Brij L23) for 5 min. Sporelings were vacuum infiltrated in the fixative for 2 min followed by another 30 min incubation period. Fixed sporelings were pelleted at 7000 g for 3 min, the fixative was removed and sporelings were washed with 1.5 ml 1x PBS. To clear sporeling tissues, fixed sporelings were incubated in 1.5 ml ClearSeeα (10% w/v xylitol, 15% w/v sodium deoxycholate, 25% w/v urea ^33^ freshly supplemented with 6.3 mg/ml sodium sulphite anhydrous^34^) in the dark for 1 week. The night before imaging, 0.2 % Renaissance SR2200 cell wall dye was added. Stained sporelings were pelleted at 7000 g for 3 min and 1.2 ml of the ClearSeeα was removed. The remaining 200 μl were transferred onto an objective slide and mounted within a gene frame. Using optimal setting, high resolution z-stacks were acquired at the Zeiss LSM 880 with a 40x/1.2 LD LCI plan-apochromat, Water, Glycerol DIC AutoCorr objective using silicone immersion-oil to closely match the refractive index of ClearSeeα^34^. The Renaissance SR2200 cell wall dye was excited by UV at 405 nm, with emission collected between 420-500 nm ^35^. For 3D reconstruction of sporelings, tiff-files of single channels (SR2200 or VENUS) were generated in ImageJ/Fiji and loaded as stacks into MorphographX ^36^ and optical cross sections were taken using the clip function. To calculate cell volumes, sporeling surfaces were blurred with the Gaussian Blur Stack-filter function (value 0.3) and cells segmented using the ITK Watershed Auto Seed function. Segmentation errors were manually corrected, and a mesh created using the Marching Cubes 3D function with a cube size of 1 and a threshold of 1500. Based on this mesh a heat map displaying cell volume was calculated using the MorphographX Heat Map function.

### QUANTIFICATION AND STATISTICAL ANALYSIS

Plasmid maps and sequence assemblies were performed in CLC Main Workbench 7.7.2 (Qiagen). Gene sequences were obtained from MarpolBase (marchantia.info). Confocal images were analysed in ImageJ/FIJI ^24^ and MorphographX ^36^. All experiments were repeated at least 2-5 times. Samples sizes (n) are indicated in the figure legends.

**Video S1**. Time lapse of prothallus formation by reproducibly oriented cell divisions, related to Figure 1.

**Video S2**. Time lapse of symmetric transverse divisions forming the plat, related to Figure 1.

**Video S3**. Time lapse of oblique divisions forming the disc, related to Figure 1.

**Video S4**. 3D reconstruction of a plate, related to Figure 1.

**Video S5**. 3D reconstruction of a disc, related to Figure 1.

**Video S6**. Time lapse of an expanding flabellum, related to Figure 2.

**Video S7**. Time lapse of notch formation on the flabellum, related to Figure 3.

**Video S8**. Time lapse of MpC3HDZ-VENUS signal at the plate to disc transition, related to Figure 5.

## Acknowledgements

We thank Katharina Jandrasits, Natalie Edelbacher and Magdalena Mosiolek (GMI) for assistance in the lab and Dr. Yasin Dagdas (GMI) for sharing the chlorsulfuron resistance cassette in Green Gate. We thank BioOptics, Molecular Biology Services, Media Lab, and Lab Support of GMI/IMBA/IMP and the VBCF Plant Sciences unit for their support. We thank Hugh Mulvey for critical reading of the manuscript. This work was funded by a grant from the Austrian Academy of Sciences to the Gregor Mendel Institute a European Research Council Advanced Grant DENOVO-P (project no. 787613) to L.D.

## Author contributions

E.-S.W. and L.D. designed the project. E.-S.W. carried out the project. E.-S.W. and L.D. wrote the manuscript.

## References

1. Goldberg, R.B., de Paiva, G., and Yadegari, R. (1994). Plant embryogenesis: zygote to seed. Science 266, 605–614. 10.1126/science.266.5185.605.

2. Strasburger, E. (1877). Ueber Befruchtung und Zelltheilung. Jenaische Zeitschrift für Naturwissenschaften 4, 435–536.

3. Aichinger, E., Kornet, N., Friedrich, T., and Laux, T. (2012). Plant stem cell niches. Annual review of plant biology 63, 615–636. 10.1146/annurev-arplant-042811-105555.

4. Leitgeb, H. (1881). Untersuchungen über die Lebermoose: Die Marchantieen und allgemeine Bemerkungen über Lebermoose. VI. (Schluss)-Heft (Leuschner & Lubensky).

5. Burgeff, H.E.N. (1943). Genetische studien an Marchantia.

6. Leitgeb, H. (1874). Untersuchungen ueber die Lebermoose (O. Deistung).

7. Shimamura, M. (2016). Marchantia polymorpha: Taxonomy, Phylogeny and Morphology of a Model System. Plant & cell physiology 57, 230–256. 10.1093/pcp/pcv192.

8. Bowman, J.L., Arteaga-Vazquez, M., Berger, F., Briginshaw, L.N., Carella, P., Aguilar-Cruz, A., Davies, K.M., Dierschke, T., Dolan, L., Dorantes-Acosta, A.E., et al. (2022). The renaissance and enlightenment of Marchantia as a model system. Plant Cell 34, 3512–3542. 10.1093/plcell/koac219.

9. Sakai, Y., Higaki, T., Ishizaki, K., Nishihama, R., Kohchi, T., and Hasezawa, S. (2022). Migration of prospindle before the first asymmetric division in germinating spore of Marchantia polymorpha. Plant Biotechnol (Tokyo) 39, 5–12. 10.5511/plantbiotechnology.21.1217b.

10. O’Hanlon, M.E. (1926). Germination of Spores and Early Stages in Development of Gametophyte of Marchantia polymorpha. Botanical Gazette 82, 215–222.

11. Kohchi, T., Yamato, K.T., Ishizaki, K., Yamaoka, S., and Nishihama, R. (2021). Development and Molecular Genetics of Marchantia polymorpha. Annual review of plant biology 72, 677–702. 10.1146/annurev-arplant-082520-094256.

12. Udar, R., and Srivastava, S.C. (1970). Sporeling Development in Preissia quadrata. Phyton, Annales Rei Botanicae, Horn 14_1_2, 165–173.

13. Jaeger, R., and Moody, L.A. (2021). A fundamental developmental transition in Physcomitrium patens is regulated by evolutionarily conserved mechanisms. Evolution & development 23, 123–136. 10.1111/ede.12376.

14. Attrill, S., and Dolan, L. (2024). Microtubules and actin filaments direct nuclear movement during the polarisation of Marchantia spore cells. bioRxiv, 2024.2002.2023.581750. 10.1101/2024.02.23.581750.

15. Menge, F. (1930). Die Entwicklung der Keimpflanzen von Marchantia polymorpha L. und Plagiochasma rupestre (Forster) Stephani. Flora oder Allgemeine Botanische Zeitung 124, 423–478. 10.1016/S0367-1615(17)32949-X.

16. Emery, J.F., Floyd, S.K., Alvarez, J., Eshed, Y., Hawker, N.P., Izhaki, A., Baum, S.F., and Bowman, J.L. (2003). Radial patterning of Arabidopsis shoots by class III HD-ZIP and KANADI genes. Current biology : CB 13, 1768–1774. 10.1016/j.cub.2003.09.035.

17. Yip, H.K., Floyd, S.K., Sakakibara, K., and Bowman, J.L. (2016). Class III HD-Zip activity coordinates lea development in Physcomitrella patens. Developmental biology 419, 184–197. 10.1016/j.ydbio.2016.01.012.

18. Prigge, M.J., Otsuga, D., Alonso, J.M., Ecker, J.R., Drews, G.N., and Clark, S.E. (2005). Class III homeodomain-leucine zipper gene family members have overlapping, antagonistic, and distinct roles in Arabidopsis development. Plant Cell 17, 61–76.

19. S.C., S., and D., S. (1995). Morphology and Sporeling Development of Fossom-bronia wondraczekii var. loitlesbergeri (Hepaticae). Phyton, Annales Rei Botanicae, Horn 35, 63–77.

20. Mehra, P.N., and Kachroo, P. (1951). Sporeling Germination Studies in Marchantiales I. Rebouliaceae. The Bryologist 54, 1–16. 10.2307/3239976.

21. Lampropoulos, A., Sutikovic, Z., Wenzl, C., Maegele, I., Lohmann, J.U., and Forner, J. (2013) GreenGate---a novel, versatile, and efficient cloning system for plant transgenesis. PloS one 8, e83043.

22. Honkanen, S., Jones, V.A.S., Morieri, G., Champion, C., Hetherington, A.J., Kelly, S., Proust, H., Saint-Marcoux, D., Prescott, H., and Dolan, L. (2016). The Mechanism Forming the Cell Surface of Tip-Growing Rooting Cells Is Conserved among Land Plants. Current biology : CB 26, 3238–3244. 10.1016/j.cub.2016.09.062.

23. Ishizaki, K., Chiyoda, S., Yamato, K.T., and Kohchi, T. (2008). Agrobacterium-mediated transformation of the haploid liverwort Marchantia polymorpha L., an emerging model for plant biology. Plant & cell physiology 49, 1084–1091. 10.1093/pcp/pcn085.

24. Schindelin, J., Arganda-Carreras, I., Frise, E., Kaynig, V., Longair, M., Pietzsch, T., Preibisch, S., Rueden, C., Saalfeld, S., Schmid, B., et al. (2012). Fiji: an open-source platform for biological-image analysis. Nat Methods 9, 676–682.

25. RStudio Team. (2020). RStudio: Integrated Development for R (RStudio, PBC.). http://www.rstudio.com/.

26. R Core Team. (2022). R: A language and environment for statistical computing (R Foundation for Statistical Computing). https://www.R-project.org/.

27. Labun, K., Montague, T.G., Krause, M., Torres Cleuren, Y.N., Tjeldnes, H., and Valen, E. (2019). CHOPCHOP v3: expanding the CRISPR web toolbox beyond genome editing. Nucleic acids research 47, W171–W174. 10.1093/nar/gkz365.

28. Chiyoda, S., Ishizaki, K., Kataoka, H., Yamato, K.T., and Kohchi, T. (2008). Direct transformation of the liverwort Marchantia polymorpha L. by particle bombardment using immature thalli developing from spores. Plant Cell Rep 27, 1467–1473. 10.1007/s00299-008-0570-5.

29. Mulvey, H., and Dolan, L. (2023). RHO GTPase of plants regulates polarized cell growth and cell division orientation during morphogenesis. Current biology : CB 33, 2897–2911 e2896. 10.1016/j.cub.2023.06.015.

30. Wallner, E.S., Dolan, L., and Bergmann, D.C. (2023). Arabidopsis stomatal lineage cells establish bipolarity and segregate differential signaling capacity to regulate stem cell potential. Developmental cell 58, 1643–1656. 10.1016/j.devcel.2023.07.024.

31. Sauret-Gueto, S., Frangedakis, E., Silvestri, L., Rebmann, M., Tomaselli, M., Markel, K., Delmans, M., West, A., Patron, N.J., and Haseloff, J. (2020). Systematic Tools for Reprogramming Plant Gene Expression in a Simple Model, Marchantia polymorpha. ACS Synth Biol 9, 864–882. 10.1021/acssynbio.9b00511.

32. Kirchhelle, C., and Moore, I. (2017). A Simple Chamber for Long-term Confocal Imaging of Root and Hypocotyl Development. Journal of visualized experiments. 10.3791/55331.

33. Kurihara, D., Mizuta, Y., Sato, Y., and Higashiyama, T. (2015). ClearSee: a rapid optical clearing reagent for whole-plant fluorescence imaging. Development (Cambridge, England) 142, 4168–4179. 10.1242/dev.127613.

34. Kurihara, D., Mizuta, Y., Nagahara, S., and Higashiyama, T. (2021). ClearSeeAlpha: Advanced Optical Clearing for Whole-Plant Imaging. Plant & cell physiology 62, 1302–1310. 10.1093/pcp/pcab033.

35. Tofanelli, R., Vijayan, A., Scholz, S., and Schneitz, K. (2019). Protocol for rapid clearing and staining of fixed Arabidopsis ovules for improved imaging by confocal laser scanning microscopy. Plant Methods 15, 120. 10.1186/s13007-019-0505-x.

36. Barbier de Reuille, P., Routier-Kierzkowska, A.L., Kierzkowski, D., Bassel, G.W., Schupbach, T., Tauriello, G., Bajpai, N., Strauss, S., Weber, A., Kiss, A., et al. (2015). MorphoGraphX: A platform for quantifying morphogenesis in 4D. Elife 4, 05864.

